# DNA-caged Nanoparticles via Electrostatic Self-Assembly

**DOI:** 10.1101/2022.11.07.515456

**Authors:** Elizabeth Jergens, Silvio de Araujo Fernandes-Junior, Yixiao Cui, Ariel Robbins, Carlos E. Castro, Michael G. Poirier, Metin N. Gurcan, Jose J. Otero, Jessica O. Winter

## Abstract

DNA-modified nanoparticles enable DNA sensing and therapeutics in nanomedicine and are also crucial for nanoparticle self-assembly with DNA-based materials. However, methods to conjugate DNA to nanoparticle surfaces are limited, inefficient, and lack control. Inspired by DNA tile nanotechnology, we demonstrate a new approach to nanoparticle modification based on electrostatic attraction between negatively charged DNA tiles and positively charged nanoparticles. This approach does not disrupt nanoparticle surfaces and leverages the programmability of DNA nanotechnology to control DNA presentation. We demonstrated this approach using a variety of nanoparticles, including polymeric micelles, polystyrene beads, gold nanoparticles, and superparamagnetic iron oxide nanoparticles with sizes ranging from 5-20 nm in diameter. DNA cage formation was confirmed through transmission electron microscopy (TEM), neutralization of zeta potential, and a series of fluorescence experiments. DNA cages present “handle” sequences that can be used for reversible target attachment or self-assembly. Handle functionality was verified in solution, at the solid-liquid interface, and inside fixed cells, corresponding to applications in biosensing, DNA microarrays, and erasable immunocytochemistry. These experiments demonstrate the versatility of the electrostatic DNA caging approach and provide a new pathway to nanoparticle modification with DNA that will empower further applications of these materials in medicine and materials science.

## Introduction

Nanoparticles modified with nucleic acids are crucial to several fields. Nucleic acid-modified nanoparticles have led to ribonucleic acid (RNA) vaccines, small interfering RNA (siRNA) therapeutics, and deoxyribonucleic acid (DNA) and RNA diagnostics in healthcare[1]. Similarly, nanoparticles modified with DNA have been used to construct complex nanostructured materials for energy applications[2]. Nanoparticle clusters[3] or arrays[4] can enable emergent behaviours, like surface plasmon resonance (SPR) or Förster Resonance Energy Transfer (FRET)[2]. DNA nanomaterials that serve as templating scaffolds for nanoparticle assembly can be created with a variety of geometries, including tiles that interlock into larger networks[5], wireframe networks and structures[6], and complex origami structures. Nanomaterials can display dynamic interactions[7, 8] that yield switchable optoelectronic properties. However, all of these applications generally require nanoparticle modification with single-stranded DNA (ssDNA) or RNA.

Creating nanoparticles modified with integrated nucleic acids to provide targeting functions can be challenging. Nanoparticle surfaces may exhibit few functional groups for modification, and even when present, different conjugation chemistries are required for each group[9]. Common conjugation approaches, such as N-hydroxysuccinimide, carbodiimide, or maleimide chemistries, have low yields[10] with high degrees of crosslinking or off-target reactions[11], and can damage nanoparticle surfaces diminishing their surface and size-dependent properties (e.g., [12]). As these approaches are not site-specific, they offer little control over the density or geometry of nucleic acid attachment. Emerging bio-orthogonal approaches with higher yields (e.g., click chemistry[10]) require the addition of unusual functional groups that still necessitate use of standard chemistries. For this reason, most DNA composite materials, including those assembled using short ssDNAs[3] or DNA nanotechnologies[2], employ gold nanoparticles that are easily modified via thiol binding to their surfaces. Very few examples exist of DNA materials incorporating other nanoparticle types (e.g., quantum dots, magnetic nanoparticles, catalysts)[13-15], despite potential advantages of such structures in biomedical imaging, optoelectronic, and energy applications. Even when considering thiol-binding chemistries used for gold nanoparticle ssDNA modification, the two main methods employed: salt aging and low pH modification require three days[3] or very precise pH control[16], which hinders their application at commercial scales. Thus, there is a critical need for universal biomodification approaches that could precisely modify broad classes of nanoparticles with nucleic acids.

One promising option for nanoparticle DNA modification is the use of DNA nanotechnology-based caging strategies[17]. DNA cages are typically constructed from interlocking tiles that form a three-dimensional network surrounding the nanoparticle(s). These systems can be simple, comprised of only a few ssDNA strands[18], or more complex systems in which many ssDNA strands first self-assemble into an intermediate structure (e.g., half of an icosahedron) that then assembles around a nanoparticle to encapsulate it (e.g., interlocking icosahedron halves)[19]. DNA cages can be designed to present ssDNA sequences on their surfaces or in their interior to controllably bind nanoparticles or attach to other DNA sequences (i.e., therapeutics, biosensing elements) or materials. However, most cage designs require *a priori* nanoparticle ssDNA modification for attachment or rely on thiol binding only applicable to gold nanoparticles[19].

Recently, Kurokawa et al.[20] demonstrated a DNA tile network assembled inside a liposome, based solely on electrostatic attraction between negatively charged DNA strands and positively charged amphiphilic headgroups in the liposome. Inspired by this example, we conceive a new strategy for nanoparticle DNA modification based on electrostatic assembly of DNA cages on the surface of positively charged nanoparticles: electrostatic DNA caging (Figure 1A). Toward universality, electrostatic DNA caging requires only a positively charged nanoparticle, does not require complicated conjugation chemistries, and does not interfere with ligands or atoms on the nanoparticle surface, preserving nanoparticle properties. In addition, cage designs include customizable ssDNA “handles” that can be used to bind external DNA sequences, such as bar-coded sensing elements or the ssDNA ends of DNA origami structures. Using ssDNA targets with partial complementary to handles enables reversible binding through the addition of ssDNA “erase” sequences with greater complementarity to handles.

**Figure 1.**
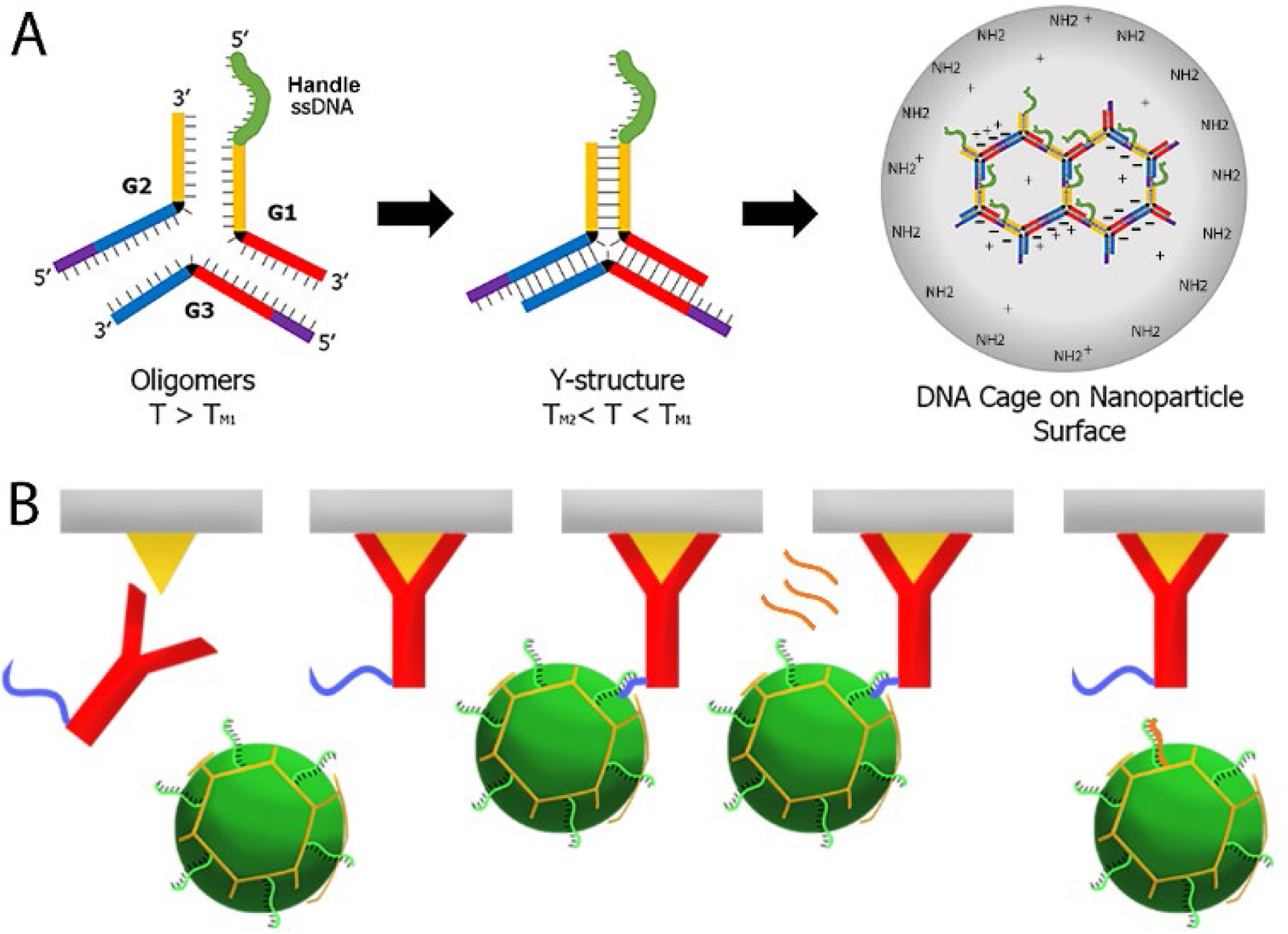
Schematics of DNA caged-NP formation and reversible sensing using ssDNA handles. (A) Single-stranded DNA (red, yellow, blue) self-assembles into Y-shaped DNA tiles that are inherently negatively-charged. DNA tiles attach to positively-charged NP surfaces through electrostatic attraction and then interlock to form an imperfect hexagonal network. Tiles are designed to present ssDNA “handle” sequences (green) extending from interlocks (purple), that are available for target binding. Reversible target binding is achieved by sequential additions of DNA with greater complementarity to handle or target sequences that initiate detachment (i.e., strand invasion). For example, in the reversible antigen labelling scheme shown in (B), antibodies pre-modified with ssDNAs bind cellular antigens through a standard immunocytochemistry approach. Then, DNA-caged nanoparticles containing fluorophores attach to ssDNA-antibodies, serving as a secondary reporter for imaging. DNA-caged nanoparticle signals are erased by adding ssDNA sequences with greater complementarity (orange) to handles, releasing DNA-caged nanoparticles from the antibody-antigen complex. Nanoparticles are then washed away to remove the fluorescence signal.

Here, we demonstrate the electrostatic DNA caging technique using a variety of nanoparticles, including polymeric micelles, polystyrene (PS) beads, superparamagnetic iron oxide nanoparticles (SPIONs), and gold nanoparticles. Using micelles as a model system, DNA cage formation was validated via direct imaging using transmission electron microscopy (TEM), indirectly by neutralization of zeta potential, and through a series of fluorescence experiments. The functionality of the handle design was then demonstrated via reversible attachment of fluorescently-labelled ssDNA sequences to DNA-caged nanoparticles in three potentially relevant media: solution phase, solid-liquid interfaces, and intracellular environments. In the latter case, a reversible immunocytochemistry labelling model was employed (Figure 1B), similar to prior examples using DNA duplexes[21]. These experiments establish electrostatic DNA caging as a flexible technology for modifying nanoparticles with DNA, compatible with a variety of healthcare and materials applications.

## Materials and Methods

### Nanoparticles

Purchased nanoparticles for DNA caging experiments were used as received. Aminated 5 nm SPIONs in aqueous suspension were purchased from Ocean Nanotech (SHA05-01, 5 mg/mL). Aminated 5 nm gold nanoparticles (0.05% gold) in aqueous suspension were purchased from Nanocs (GP5-AM-1). PS beads with amine (PS20-AM-1), carboxylic acid (PS20-CA-1), or hydroxy (PS01-20-1) functionalization were purchased from Nanocs. All PS beads were 20 nm in diameter and at 0.01% by volume in an aqueous solution.

### Synthesis and purification of micelle nanoparticles

In addition to purchased nanoparticles, lipid-polymer micelles were used as a model system. Micelles[22, 23] encapsulating coumarin-6 dyes for visualization were prepared using electrohydrodynamic mixing-nanoprecipitation (EM-NP) as we described previously[24]. Briefly, 1,2-Distearoyl-sn-glycero-3-phosphoethanolamine-N-[methoxy(polyethylene glycol)-2000] (DSPE-PEG, Avanti Polar Lipids Inc. cat. no. 88010P) and 1,2-Distearoyl-sn-glycero-3-phosphoethanolamine-N-[methoxy (poly-ethylene glycol)-2000]-NH_2_ (DSPE-PEG-NH2, Avanti Polar Lipids Inc. cat. no. 88128P) were each dissolved in tetrahydrofuran (THF) with butylated hydroxytoluene free radical inhibitor (Sigma-Aldrich cat. no. 360589) at a concentration of 5 mg/mL. For 100% NH2 experiments only DSPE-PEG-NH2 was used, whereas for 50% NH2 experiments these were mixed in equal proportions. Solutions were heated for 30 sec in a 37 °C water bath to promote dissolution and vortexed for 10 sec. Separately, coumarin-6 dye (1 mg/mL in THF, purity ≥ 99%, Sigma-Aldrich cat. no. 546283) was prepared by sonicating for 1 min and vortexing for 10 sec. Particles were generated by mixing 200 mL polymer solution, 10 mL coumarin-6, and 190 mL THF and spraying 200 mL of this solution into 10 mL of distilled water at a flow rate of 12.7 mL/hr, under a voltage of -2500 V, for 45 sec. To ensure the purity of the water in which the micelles were prepared, we observed an acceptable maximum initial electrical current rate of 0.29 mA and a final maximum of 1 mA. The resultant nanocomposites were purified via centrifugal filtration (regenerated cellulose 30 kDa NMWL Amicon Ultra-15, Millipore Sigma cat. no. UFC903024) at 4000 rpm for 15 minutes with 3 washes using DI water. Resultant nanocomposites had a coumarin-6 encapsulation efficiency of ∼ 30-50% with sizes ranging from 20-30 nm (Supplementary Methods, Supplementary Figures 1-2). Nanocomposites used in fluorescence saturation studies were manufactured via the same method with equal amounts of DSPE-PEG-NH2 and DSPE-PEG-Cy3 (Nanosoft Polymers cat. no. 4586) (i.e., no DSPE-PEG) without coumarin-6. [Note that in these experiments Cy3 is on the micelle surface, in contrast to hydrophobic coumarin-6 dyes encapsulated in the micelle interior.]

### Formation of DNA nanostructures and DNA cages

DNA cages were assembled from G1, G2, and G3 sequences (Supplementary Table 1) to form three-way junction, Y-shaped DNA tiles as described by Kurokawa et. al.[20] with modifications (Figure 1A). These structures are designed to form a network on curved surfaces. Here, all strands were modified with “handle” strands (Figure 1A, green, AAAAATTTCGACGTTACATGCACCTC) extending from sticky end interlocking strands (Figure 1A, purple). Also, for surface and interlock quenching experiments, 1/120^th^ of G3 strands were modified with black hole quencher (BHQ-1) on their sticky ends. For surface saturation and interlock quenching experiments, 1/120^th^ of the G2 strands were conjugated to FAM-6 (fluorescein).

Custom DNA oligos were purchased from Integrated DNA Technologies (IDT) as lyophilized powders. Each oligo was reconstituted in 20 mM Trizma base (Sigma-Aldrich cat. no. T6066) with 350 mM NaCl (Sigma-Aldrich cat. no. S7653). A solution of G1, G2, and G3 strands, each at 18 μM, was heated at 80 °C for 10 minutes and slowly cooled to 4 °C to form Y-shaped tiles. Tiles were stored at 4 °C until use. DNA caged nanoparticles were formed by incubating DNA with nanoparticles at specified ratios (DNA: PEG molar ratios of 2-18, Table 1) at room temperature for 10 minutes. Samples were purified using centrifugal filtration (regenerated cellulose 100 kDa NMWL Amicon Ultra-15, Millipore Sigma cat. no. UFC910024) at 4000 rpm for 15 minutes with 3 washes of phosphate buttered saline (PBS, ThermoFisher cat. no. 28372). After purification, DNA-caged nanoparticles were used immediately.

**Table 1.**
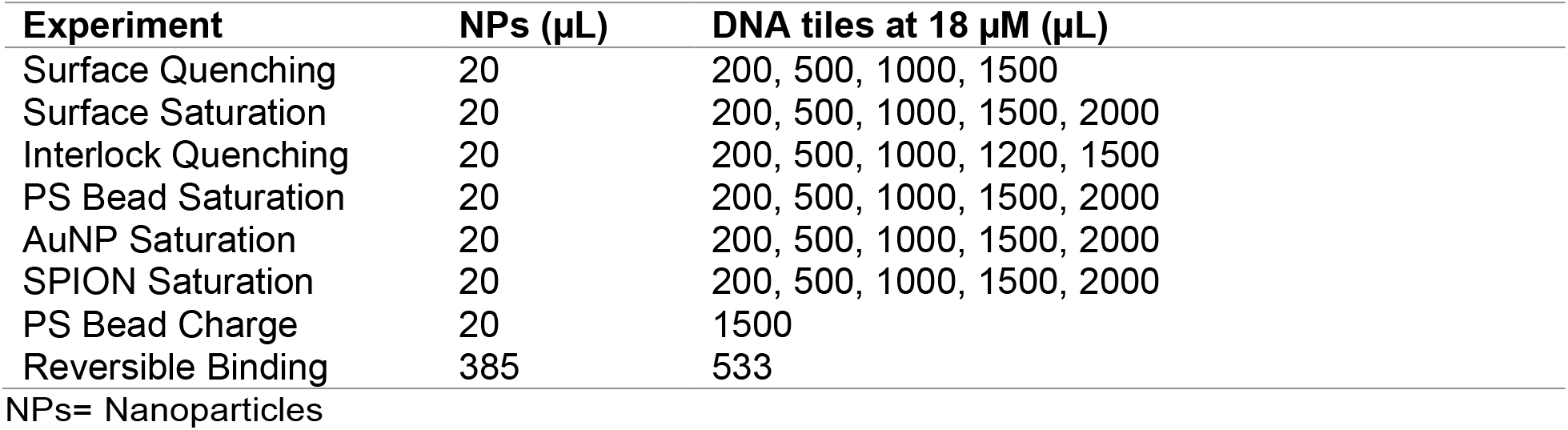
Experimental conditions for DNA tile adsorption to nanoparticles.

### Characterization of DNA-caged nanoparticle materials

Nanoparticle and DNA-caged nanoparticle sizes were characterized using dynamic light scattering (DLS) (BI 200-SM, Brookhaven Instruments Corp.). All measurements were performed with a 90-degree collection angle with the aperture set to 200x. Selected samples were also imaged using transmission electron microscopy (TEM) to confirm morphology. TEM samples were prepared on copper, square mesh grids (400 mesh, Electron Microscopy Services cat. no. CF400-CU) that were cleaned and hydrophilized using a PELCO easiGlow™ Glow Discharge Cleaning System. Samples were deposited on grids and negatively stained with 1% uranyl acetate. Images were collected with an FEI Tecnai G2 Bio Twin TEM at 60-75 eV. Neutralization of nanoparticle surface charge by DNA cage adsorption was assessed using zeta potential (ZP) with a Brookhaven Instruments Corp. ZetaPALS.

DNA cage formation was characterized through a series of fluorescence experiments, employing micelles manufactured as described above but without coumarin-6 to avoid interference. In surface quenching experiments, BHQ-1-DNA tiles were incubated with fluorescent micelles displaying Cy3 molecules on their surfaces, whose fluorescence was quenched with tile adsorption. In surface saturation experiments, FAM-6-tiles were incubated with aminated micelles, whose fluorescence increased with tile adsorption. In interlock quenching experiments, BHQ-1- and FAM-6-DNA tiles were incubated with aminated micelles, whose fluorescence increases and then quenches to indicate tile adsorption and interlocking, respectively. All experiments were performed at DNA: PEG molar ratios of 2-18 at the volumes and concentrations listed in Table 1 as described above. Following incubation, samples were purified using centrifugal filtration (regenerated cellulose 100 kDa NMWL Amicon Ultra-15, Millipore Sigma cat. no. UFC910024) at 4000 rpm for 15 minutes with 3 washes of phosphate buttered saline (PBS, ThermoFisher cat. no. 28372). Fluorescence of DNA-caged nanoparticles was then measured using a Photon Technology International (PTI)-810 fluorometer.

### Solution-phase reversible ssDNA binding to DNA-caged nanoparticles

To confirm functionality of ssDNA cage handles, reversible ssDNA binding of a fluorescence reporter system was employed. DNA cages for these experiments were prepared as described above using the values shown in Table 1 in PBS. Fluorescently labelled ssDNA sequences with partial complementarity to ssDNA handles were added to coumarin-6-DNA-caged nanoparticles, permitting their attachment via hybridization. These could then be removed by adding non-fluorescent ssDNA “erase” sequences with increasing binding complementarity, verifying handle functionality. “Target” Cy5-ssDNA sequences with 9 complimentary base pairs (bps) (TAAATTGAGGATTATCAAA**CATGTAACG**/3Cy5Sp/) were added to DNA-caged nanoparticles at ∼ 650 times molar excess in water to the DNA tiles used (in PBS). Samples were incubated for 15 minutes before excess Cy5-ssDNA was removed via centrifugal filtration as described in the cage formation methods above. The fluorescence of the bound Cy5-labeled target strands was then measured via spectrofluorometry as above. This process was repeated with three additional strands of increasing complementarity, alternating between fluorescent and non-fluorescent ssDNAs: non-fluorescent ssDNA with 12 complimentary bps (ATTGAGGATTATCAAA **GAGGTGCATGTA**), Cy-5 ssDNA with 15 complimentary bps (GATTATCAAA**GAGGTGCATGTAACG**/3Cy5Sp/), and non-fluorescent ssDNA with 26 complimentary bps (**GAGGTGCATGTAACGTCGAAATTTTT**, full complement). As a control, we also employed non-complementary (/5Cy5Sp/TTTTTTTTTTTTT/3ThioMC3-D/). To demonstrate compatibility with biological reagents, this experiment was repeated in detergent buffers. DNA-caged nanoparticles were incubated in 0.25% Tween-20 (Sigma-Aldrich cat. No. P1379) in PBS or 0.25% Triton X-100 (Sigma-Aldrich cat. No. X100) in PBS for 1 hr. Then, repeated binding cycles were performed as described above.

### Reversible DNA-cage binding at solid-liquid interfaces

To demonstrate reversible DNA-caged nanoparticle binding to ssDNA at the solid-liquid interface, we employed ssDNA bound to microscope slides, similar to the presentation of DNA microarrays. Standard microscope cover glass (24 × 60, Fisher Scientific cat. no. 12-548-5P) was coated with 10 nm of gold using a thin film deposition system (Kurt J. Lesker Co. Lab-18). Then, 80 µL of thiolated DNA (GATTATCAAA GAGGTGCATGTAACGTCG/3ThioMC3-D/) was reduced with 20 µL of 500 mM DL-Dithiothreitol (DTT, Sigma-Aldrich cat. no. 43815) in 500 mM NaH2PO4 (Sigma-Aldrich cat. no. 71505) and 500 mM Na2HPO4 (Sigma-Aldrich cat. no. S5136) (pH=8.4). This mixture was placed on a vortex mixer for 1 hr at 1000 rpm. Concurrently, a NAP-10 column (G-25 DNA grade, Cytiva cat. no. 17085401) was equilibrated with 5 mL DI water passed through the column three times. After vortexing, the DNA sample and 400 µL of DI water were added to the column, and the first fraction was collected. Then, 500 µL of DI water was added to produce a second fraction. This was repeated twice more. The third fraction contains a majority of the reduced ssDNA, with DTT removed. The reduced ssDNA was then introduced to gold-coated slides and incubated overnight. Samples were then salt-aged by adding 12.5 µL of 1 M NaCl (Sigma-Aldrich cat. no. S7653) and 0.1 M sodium phosphate (pH=7) each hour for 4 hr. The slides were incubated overnight, then washed with PBS (ThermoFisher cat. no. 28372).

To confirm handle functionality, coumarin-6-DNA-caged nanoparticles were prepared as described above using values shown in Table 1. A droplet (2 µL) of DNA-caged nanoparticles was placed on the ssDNA-coated slide surface and allowed to attach for 15 minutes. The droplet was then imaged on an inverted fluorescent microscope (Nikon, IX81) at 4x magnification before removal by aspiration to assess the maximum fluorescence (bound and unbound nanoparticles). The slide was then washed with PBS three times and imaged again (under a droplet of PBS for appropriate refraction) to identify the bound fraction of DNA-caged nanoparticles. To remove bound DNA-caged nanoparticles from the slide surface, 2 µL of ssDNA (20 nM) with 26 complimentary bps (GAGGTGCATGTAACGTCGAAATTTTT, full complement) was added and incubated for 15 minutes. The droplet was then removed, and the slides were washed three times with PBS. The sample was then imaged (under PBS) to identify the “erased” fraction of ssDNA-caged nanoparticles. The negative control was conducted following the same method using a slide without ssDNA to identify non-specific attachment.

### DNA binding inside fixed cells: reversible immunocytochemistry

To establish compatibility with bioassays, reversible DNA-caged nanoparticle binding was assayed using an *in vitro* immunocytochemistry model. Actin cytoskeletal protein, a cytoplasmic protein expressed in every cell, was labelled to provide an easy assessment. In this approach (Figure 1B), DNA-conjugated antibodies were used to recognize antigens. Then, coumarin-6-DNA-caged nanoparticles presenting ssDNA partially complementary to the ssDNA-antibodies were used as secondary labelling reagents. After fluorescence imaging, DNA-caged nanoparticles were detached from ssDNA-antibodies using erase sequences with increasing complementarity, as described above.

### DNA-Antibody conjugation and characterization

Primary actin antibodies (β-Actin Monoclonal Antibody (AC-15), Life Technologies AM4302) were modified with azide groups using GlyCLICK azide activation kits (Genovis L1-AZ1) before use in a copper-free click chemistry reaction. GlyCLICK permits site selective antibody modification of up to 4 sugars present on the heavy chains, precluding modification of the antigen binding pocket. Amine-terminated Cy5 (to verify attachment)-ssDNA (/5Cy5/GATTATCAAAGAGGTGCATGTAACGTCG/3AmMO/, 18 complementary bps) sequences were modified with dibenzocyclooctyne (DBCO) for click chemistry reaction with antibody azides by incubation with dibenzocyclooctyne (DBCO)-sulfo-N-Hydroxysuccinimide (NHS)-ester (Click Chemistry Tools cat. no. A124) at a 1:1000 molar ratio for 30 min at 37 °C. DBCO-DNA was then mixed at 20 times molar excess to azide-activated antibodies for 2 hrs at room temperature. After the click reaction, ssDNA-antibodies were purified using centrifugal filtration at 5000 x g for 6 minutes (Amicon Ultra-0.5 cat. no. UFC5050) using columns pre-equilibrated with 500 µL at the same settings. DNA conjugation efficiency was quantified by measuring DNA absorbance using UV-vis spectroscopy (Fisher Scientific Genesys 6) and antibody concentration using a micro BCA protein assay kit (ThermoFisher Scientific cat. no. 23235), and was ∼80% with 1-2 DNA strands per antibody.

### Reversible cell labelling

U87-MG (ATCC cat. no. HTB-14) human glioblastoma cells were cultured in Dulbecco’s modified Eagle medium/nutrient mixture F-12 (DMEM/F-12, ThermoFisher Scientific cat. no. 11330057) supplemented with 10% fetal bovine serum (FBS premium grade, VWR cat. no. 97068-085), 1% penicillin-streptomycin (Fisher Scientific cat. no. 15140122), and 1x MycoZap (VWR cat. no. NC9023832) following the manufacturer’s instructions. Prior to use, cells were cultured at 37 °C with 5% CO2 and passaged at 80% confluency. Cells (100k) were then seeded in a single chamber transmission flow cell (FC 81-AL, Biosurface Technologies corp.) in 2 mL growth media and incubated overnight (∼18 hr). The next day, cells were washed with PBS and fixed with 4 wt.% paraformaldehyde (PFA, Sigma Aldrich cat. no. 158127) for 20 min at room temperature (RT). Cells were washed three times with PBS for 5 min each, and the chamber was imaged on an Olympus IX81 fluorescence microscope. Liquid addition or removal were performed at a flow rate of 10 mL/hr; incubation steps (i.e., stop-flow) were performed under no flow.

First, cells were permeabilized with 0.25% Triton in PBS for 30 min at room temperature. Then, cells were washed with 5 mL of PBS and incubated with a blocking buffer comprised of 1% bovine serum albumin (BSA, Fisher Scientific cat. no. BP9703) and 10% goat serum (Sigma Aldrich cat. no. G9023) in PBS for 2 hr at room temperature. In a standard experiment, ssDNA-anti-β-actin primary antibody (ThermoFisher Scientific cat. no. AM4302) was added to cells at 1:200 dilution in blocking buffer and incubated overnight at 4 °C. As a positive control, anti-β-actin primary antibodies without ssDNA modification were also employed. Cells were then washed with 5 mL of PBS. For experimental samples, DNA-caged nanoparticles were introduced to the flow cell at a concentration of 1000 μg/mL and incubated for 1 hr in the dark at room temperature. For positive control experiments, Alexa-568 goat anti-mouse secondary antibodies (Abcam cat. no. 175473) were flowed into the cell at a 1:200 dilution in blocking buffer and incubated for 1 hr in the dark at RT. There were two positive control experiments, employing either unmodified or ssDNA modified anti-β-actin primary antibodies. In all experiments, cells were then washed with 5 mL of PBS and counter-stained with DAPI (Sigma Aldrich cat. no. D9542) for 5 min. A final wash of 5 mL PBS was performed before observation. Samples were imaged at 20X using MetaMorph software, with positive control samples imaged using the TRITC channel (excitation: 540-560, emission: 570-640) and DNA-caged nanoparticle samples imaged using the FITC channel (excitation: 460-500, emission: 520-560). DAPI signal was imaged using the DAPI channel (excitation: 330-370, emission:440-490). To erase the DNA-caged nanoparticle fluorescent signal, erase ssDNA (26 bps complimentary to the handle strand) (20 nM in PBS) was flowed into the cell and incubated under stop-flow for 15 minutes. (This removes DNA-caged nanoparticles but does not disrupt antibody-antigen binding). Time-lapse images were collected at 30-second intervals while the erased DNA was flowed in for 30 min after. (DAPI signal was not imaged in timelapse because time constraints limited switching between imaging channels.) After the 15 min incubation step, the first wash with 5 mL of PBS was conducted. After 10 minutes, a second wash with 5 mL of PBS was conducted. After 10 more min, a third wash with 5 mL of PBS was conducted. Images were captured as described above to show erase, then another solution of erase DNA was added as stated above. Following this, cells were washed with 5 mL PBS and a second label/erase cycle was performed. However, no additional antibodies were added as the first cycle erased only DNA-caged nanoparticles and not primary antibodies.

### Image Analysis

The fluorescent intensity of three individual cells from three different sites for a total of 9 cells was measured as the raw integrated grey value using ImageJ (NIH). The background signal was subtracted from the raw values in each cell before being normalized to the initial signal intensity. Any frames with obvious dust or other contaminants were excluded from the image analysis.

### Statistical Analysis

All statistical analysis was conducted using JMP 16 statistical analysis software at a significance level of 0.05. First, an ANOVA was performed to confirm the presence of a statistically significant difference between the values observed. Then, Tukey tests were performed to determine statistical significance of individual pairings.

## Results

### Verifying DNA Cage Formation on Nanoparticle Surfaces

DNA oligomers of alternating sequences were designed to interlock with each other and used to form a cage-like structure on the surface of nanoparticles (Figure 1A). This design, based on DNA tiles used to form artificial cytoskeleton inside liposomes[20], relies on electrostatic attraction between positively charged nanoparticles and negatively charged DNA sequences for self-assembly. The cage is stabilized by the formation of interlocks between tiles, preventing desorption. As a model system, most of our experiments employed block copolymer micelles formed using electrohydrodynamic mixing[25]. We verified cage formation using TEM and a series of fluorescence labelling and quenching experiments.

First, we evaluated DNA-caged nanoparticle morphology using TEM with negative staining compared to nanoparticles alone and self-assembled DNA tile nanostructures alone. DNA nanostructures alone appeared as dark spheres ∼30-40 nm in diameter. In contrast, micellar nanoparticles appeared as white circles ∼10-20 nm in diameter (Figure 2A, B). These results were both consistent with prior reports[18, 25]. DNA-caged nanoparticles were ∼20-30 nm in diameter, closer in size to the larger DNA nanostructures than polymer micelles, but their appearance in TEM was distinct. Structures displayed dark regions consistent with DNA nanostructures, surrounded by a lighter halo more closely resembling polymer micelles (Figure 2C). These data suggest formation of a unique structure differing from either DNA nanostructures or micelles.

**Figure 2.**
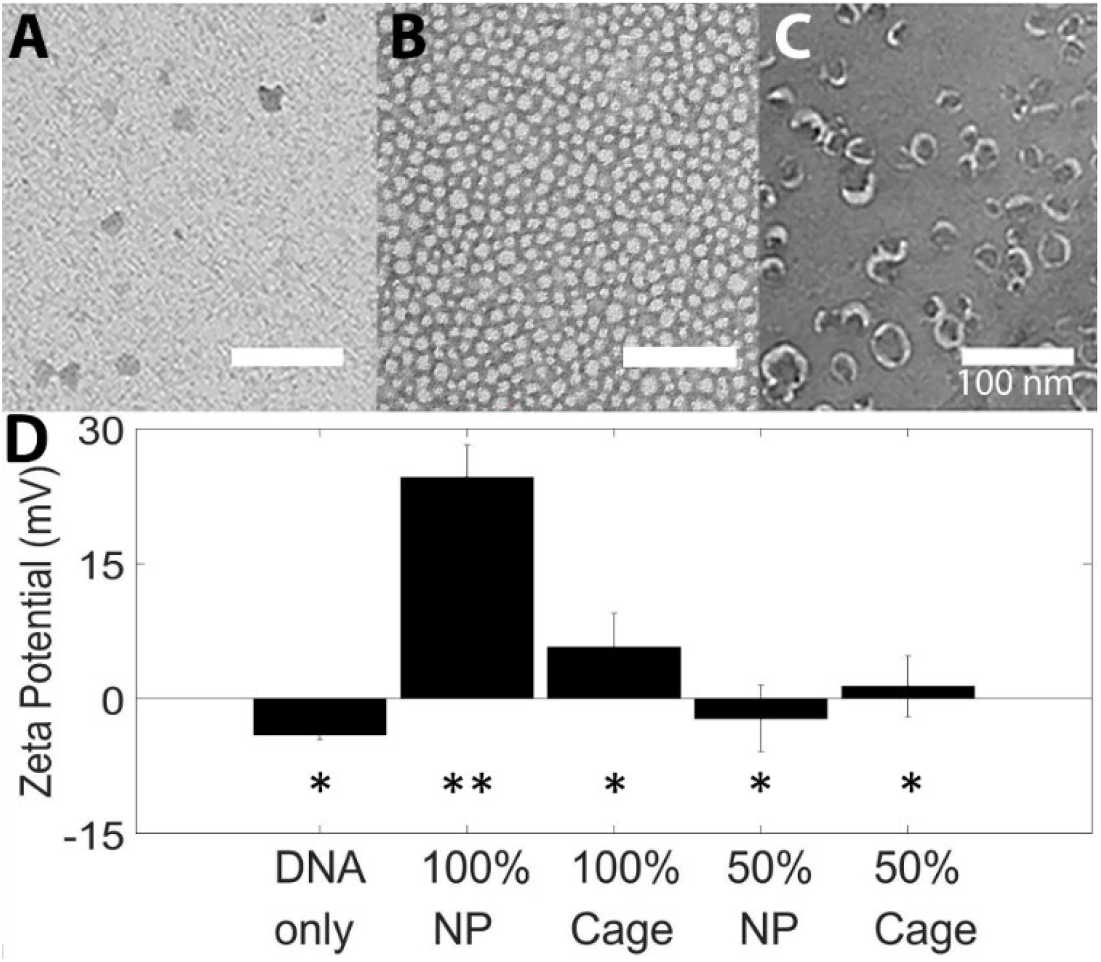
Formation of DNA-caged nanoparticles. A-C) Negatively stained TEM images of A) DNA nanostructures, B) nanoparticles, and C) DNA-caged nanoparticles. Scale bar is 100 nm. D) Zeta potentials of: DNA nanostructures, nanoparticles with 100% (100 NP) or 50% (50 NP) NH_2_ terminated block copolymers, and DNA-caged nanoparticles with 100% (100 cage) and 50% (50 cage) NH_2_ terminated block copolymers. Differing numbers of asterisks indicate statistical differences between samples.

TEM results were corroborated by evaluating zeta potential of DNA nanostructures, nanoparticles, and DNA-caged nanoparticles (Figure 2D). Nanoparticle micelles for these experiments were formed from block copolymers with either 100% positively-charged terminal amine groups or 50% amine and 50% neutral, hydroxyl terminal groups. Consistent with negative charges associated with the DNA phosphate back bone, DNA nanostructures alone displayed a slightly negative zeta potential. Uncaged 100% NH_2_ nanoparticles displayed the expected positive zeta potential consistent with their terminal amine groups, whereas 50% NH_2_ nanoparticles had zeta potentials of ∼ 0. This may reflect burying of terminal NH_2_ groups in the micelle core[26], which would reduce surface charge. After the caging process, the zeta potential of 100% NH_2_ DNA-caged nanoparticles was reduced and closer to zero, consistent with the expected electrostatic adsorption of DNA tiles on the micelle surfaces. Zeta potential of 50% NH_2_ DNA-caged nanoparticles was unchanged (not statistically different). It has been reported that non-specific binding of nanoparticles to the cell plasma membrane can be minimized by a neutral zeta potential[27]. To reduce non-specific binding, subsequent experiments were performed with micelles comprised of block copolymers with 50% NH_2_ termination that displayed zeta potentials close to zero.

These data establish the formation of a unique composite structure that includes micelle nanoparticles and DNA tiles. To further characterize DNA tile electrostatic adsorption and interlocking on the micelle surface, we employed a series of fluorescence and fluorescence quenching experiments (Figure 3, Supplementary Figure 3). The first experiment employed micelles comprised of block copolymers with terminal Cy3 fluorophores whose fluorescence could be quenched by adsorption of DNA tiles modified with black hole quencher-1 (BHQ-1)[28] (Figure 3A). DNA tiles were added to polymer micelles at DNA to PEG molar ratios of 2-18. Signal decreased with increasing addition of tiles; and at ratios higher than 14:1, fluorescence decreased by 95% from the initial signal. These results imply electrostatic adsorption of DNA tiles to the micelle surface, consistent with zeta potential results.

**Figure 3.**
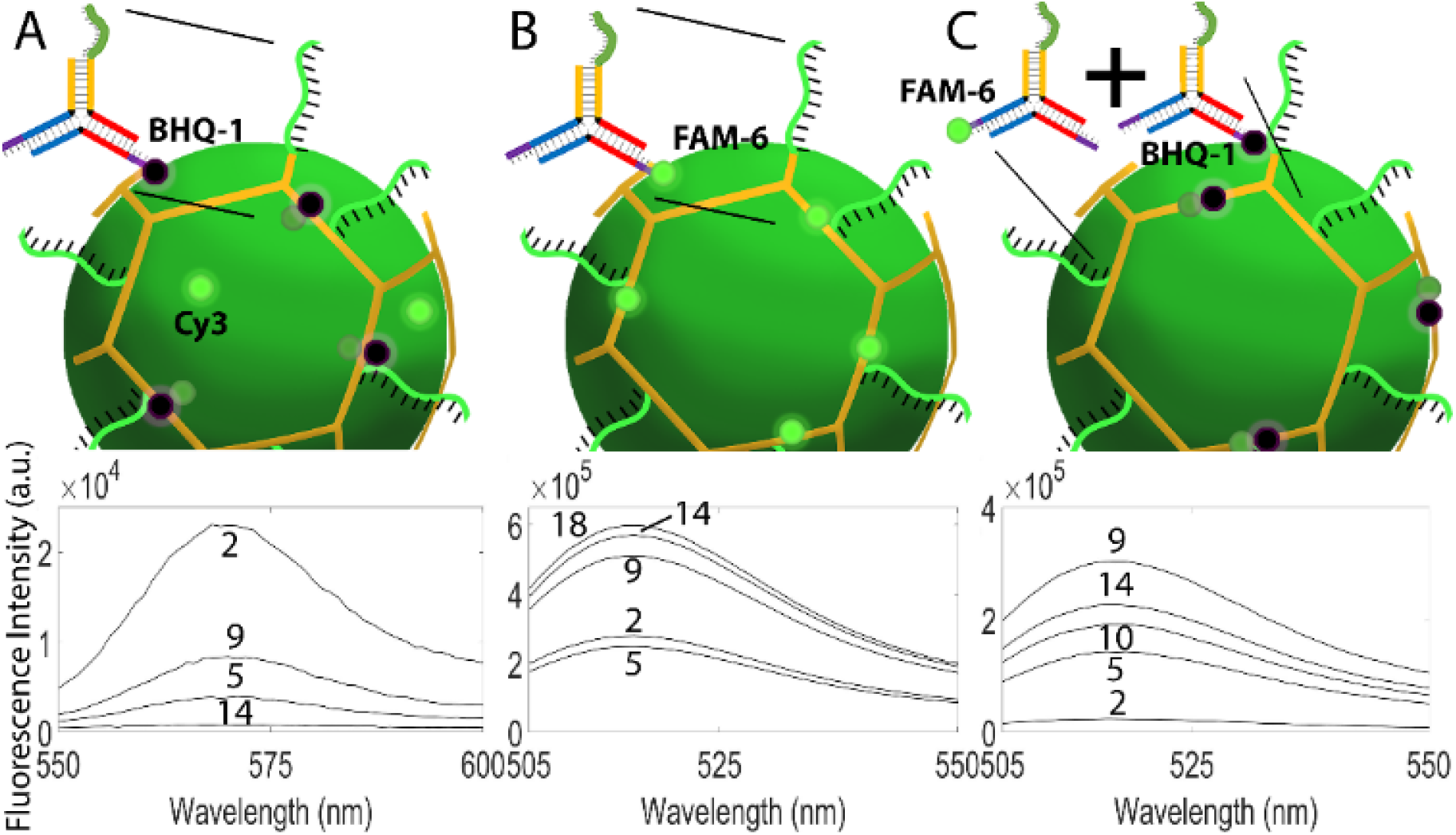
DNA tile cage formation and interlocking were confirmed by fluorescence experiments. A) Electrostatic adsorption of black hole quencher (BHQ-1) labelled DNA tiles to surfaces of fluorescent Cy3-micelles was confirmed through fluorescence quenching, indicating tile attachment. Fluorescence intensity for molar ratios of 2-14 is shown. The signal was reduced by 95% at the highest molar ratios examined. B) Saturation of tiles on the micelle surface was identified by adsorption of DNA tiles modified with FAM-6 dye. Fluorescence intensity for molar ratios of 2-18 is shown. Saturation was observed at DNA: polymer molar ratios of ∼14-18. C) Tile interlocking was confirmed by fluorescence quenching between strands on interlocking DNA tiles presenting FAM-6 dye and BHQ-1. Fluorescence intensity for molar ratios of 2-14 is shown. Curves are labelled with DNA: PEG molar ratio. Figures show representative curves for selected molar ratios, averages across replicates can be found in Supplementary Figure 3.

Next, adsorption of DNA tiles and saturation of micelle surfaces was evaluated using DNA tiles modified with FAM-6 fluorophores (Figure 3B). Tiles were added to micelles at increasing molar ratios of 2-18. Fluorescence of the purified micelles was then assessed to evaluate tile binding. Fluorescence increased with tile addition; however, increases in fluorescence diminished with increasing molar ratio. At a molar ratio of ∼14:1, a saturation limit is reached after which the further addition of DNA tiles does not result in a significant increase in micelle fluorescence. The signal at this ratio is about 2-3 times greater than that of the lowest ratio (2:1). These results further confirm DNA tile electrostatic adsorption to the micelle surface, and identify saturation limits in this system, which appears to follow a Langmuir behaviour. However, it is unlikely that the surface is completely covered with DNA tiles. At high molar ratios, steric hindrance of bound DNA tiles likely limits subsequent tile binding. These experiments provide a method for identifying optimal adsorption conditions, but do not prove the existence of an interlocked DNA tile-based cage.

To confirm the formation of interlocks, a fluorescence quenching strategy was employed in which DNA tiles presenting FAM-6 fluorophores on strands complementary to tiles presenting BHQ-1 were used (Figure 3C). DNA tiles were added to micelles at molar ratios of 2-18. In these experiments, fluorescence increased with DNA tile addition up to molar ratios of 9. This likely reflects a low-density regime in which tiles are too dilute to form interlocks. At molar ratios of 10, fluorescence decreases by 37%, indicating transition to a regime in which interlocking can take place. Initially, DNA tiles are randomly adsorbed to the nanoparticle surface. As density increases, the structures begin to interlock. However, at higher molar ratios, fluorescence increases again. In this regime, steric hindrance becomes an issue, and interlock formation is reduced. Additionally, the hexagonal nature of these DNA tiles does not allow for the complete formation of a sphere as stated by Euler’s formula *V* − *E* + *F* = 2, where V is the number of vertices, E is the number of edges, and F is the number of faces[29]. A sphere comprised of hexagons must include at least 12 pentagons. Thus, these DNA tiles will not form perfect interlocking structures, but this is not necessary, as the combination of electrostatic interactions and physical interlocks stabilizes the structure. Other DNA cage designs based on DNA tiles, such as those used to form icosahedra[19], have to potential to completely inscribe a sphere, and may be a good alternative for a completely interlocked structure.

These data establish DNA cage formation on micelle surfaces with at least partial interlocking. Next, the versatility of this approach was demonstrated by forming DNA-caged nanostructures using polystyrene (PS) beads, gold nanoparticles, and SPIONs. For each nanoparticle type, a saturation curve was generated as in Figure 3B (Supplementary Figure 4), indicating DNA tile adsorption to nanoparticle surfaces. In addition, PS beads with different surface charges were evaluated to confirm electrostatic attraction as the main method for DNA cage assembly. Neutral and negatively-charged PS beads showed less DNA binding than positively-charged PS beads (Supplementary Figure 5); however, some tile binding was still observed. This likely reflects non-specific binding as values were unchanged between -OH and -COOH terminated PS beads. The statistically significant increase in binding for NH_2_ modified beads indicates electrostatic tile adsorption.

### Solution-Based Reversible DNA Binding

DNA-modified nanoparticles are often studied in solution phase, for example as aggregation-based biosensors[30] or as components of DNA-based nanomaterials assembled through base-pairing interactions[2]. In these experiments, the functionality of DNA cage “handles” (i.e., green sequences in Figure 1A) was investigated. Handle sequences could be inaccessible for a variety of reasons, including steric effects[31] and adsorption to the nanoparticle surface[32]. Handle functionality was evaluated using strand displacement reactions to alternately bind fluorescent and non-fluorescent ssDNA sequences to DNA cage handles. Strand displacement reaction rate is dependent upon the length, concentration, and composition of the ssDNA[33], permitting strands of increasing length and complementarity to displace shorter strands. These experiments employed polymer micelles with DNA cages formed at an 18:1 DNA:polymer ratio with a 26 bp handle. DNA-caged nanoparticles were first exposed to an excess amount of Cy5-labeled 9 bp ssDNA, and binding was confirmed via fluorescence. Strands were erased by adding ssDNA sequences with greater handle complementarity lacking Cy5 fluorophores (Figure 4A). This process was repeated with strands of increasing complementarity (i.e., 15 bp fluorescent Cy5-ssDNA and 26 bp erase ssDNA) for a second cycle.

**Figure 4.**
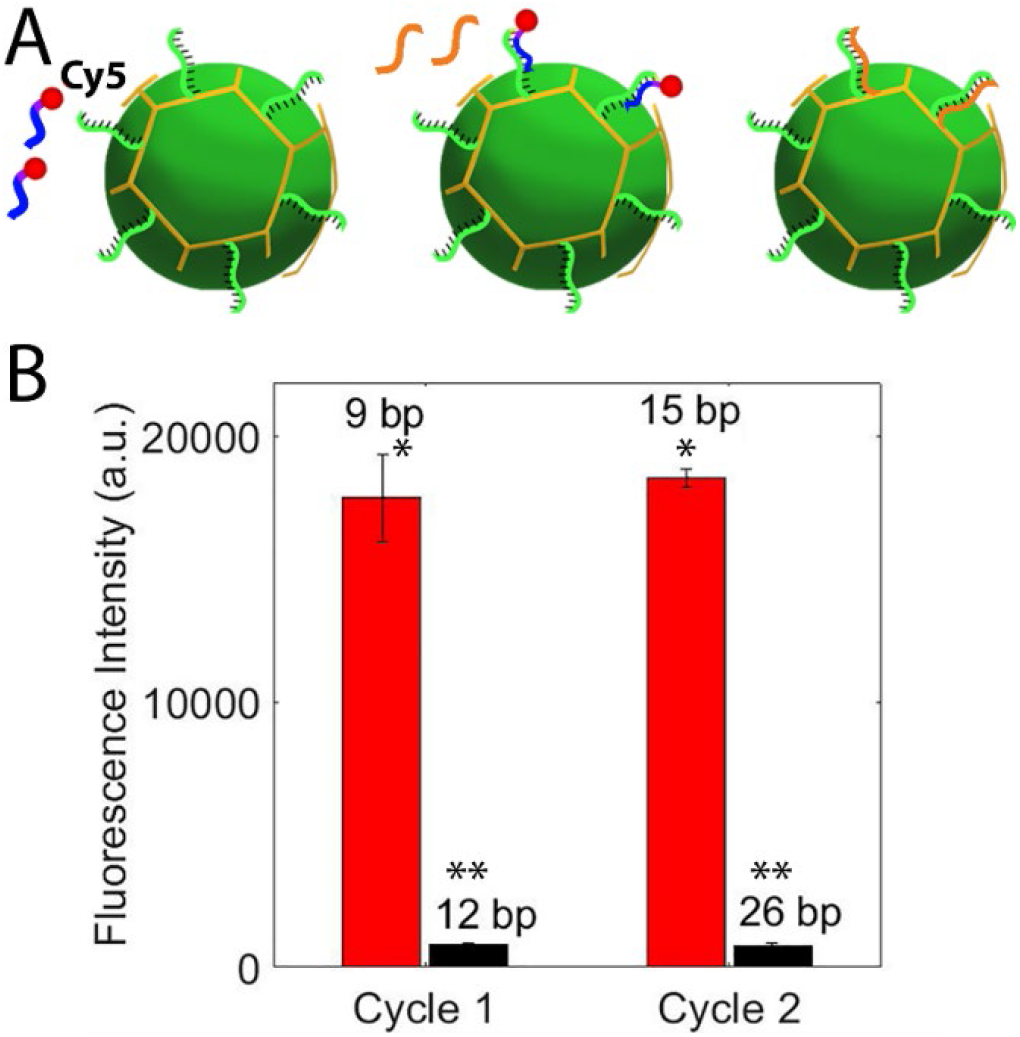
Reversible binding of fluorescent ssDNA sequences to DNA cage ssDNA handles in solution. A) Schematic showing DNA caged nanoparticle presenting 26 base pair (bp) ssDNA handle sequences (green) for binding. Fluorescent, Cy5 (red)-ssDNA (blue) sequences with partial complementarity to handles are added and fluorescence recorded. Then, erase ssDNA sequences (orange) with greater handle complementarity are used to remove Cy5-labeled strands. B) Fluorescent intensity of DNA-caged nanoparticles after Cy5-ssDNA binding and erase over two cycles. Values listed on the graph are the number of complimentary bps in each strand. Differing numbers of asterisks indicate statistical differences between samples.

Each cycle shows an ∼ 95% decrease in fluorescence from the initial fluorescence signal (Figure 4B), suggesting reversibility of ssDNA binding to DNA cage handles. Very little difference in signal is evidenced between cycles, suggesting that the 5% residual may reflect non-specifically bound or sterically trapped DNA strands. Further, the fluorescence signal intensity is retained between cycles, indicating that handles remain available for binding. In contrast, the fluorescence signal of samples incubated with non-complimentary DNA (3700 ± 440) was statistically different from both labelled and erased samples. We also evaluated handle functionality in the presence of two surfactants, Tween-20 and Triton X-100, commonly used in biological protocols to improve cell or tissue permeability. These surfactants can disrupt polymer micelles; thus, these experiments indirectly evaluate DNA cage stability. As in PBS, strand displacement could be repeated with high efficiency (Supplementary Figure 6). It is likely that erase efficiency could be increased by optimizing erase ssDNA incubation times and concentrations, which was not investigated in this work. Given this successful validation, strand displacement was used in subsequent experiments to show reversible ssDNA binding.

### Reversible DNA caged nanoparticle binding to surfaces

Another common presentation for DNA-modified nanoparticles is attachment to surfaces, for example as they might be used in gene arrays. To evaluate handle functionality and reversible binding at solid-liquid interfaces, we evaluated the ability of DNA caged nanoparticles encapsulating coumarin-6 dye to reversibly attach to DNA labelled slides. Fluorescence of a liquid droplet containing DNA-caged nanoparticles was measured before, during, and after DNA cage attachment (Figure 5, Supplementary Figure 7). Whereas a large fluorescence increase was seen after deposition of the droplet on the slide surface (i.e., PBS blank vs. DNA cage droplet), fluorescence remaining after droplet removal and washing (i.e., attached DNA cages) increased by only ∼40% from the blank sample. This increase was higher than and statistically significant from non-specific attachment, which was only ∼10% higher than that of the blank sample. Signal following erase was lower and statistically different from the signal of bound DNA cages, and was not statistically different from the non-specific attachment signal. These data suggest that DNA caged nanoparticles remaining post-erase are most likely non-specifically adsorbed to the slide surface. The significant decline in fluorescence between the droplet and attached DNA cages most likely results from a combination of low cage diffusion to the slide surface and potential saturation of slide ssDNA binding sites. Importantly, signal declines after erase indicate that most DNA-caged nanoparticles could be erased from slide surfaces. The remaining signal represents the non-specifically bound portion of DNA-caged nanoparticles. Non-specific attachment was low (only 10% increase), but could be further reduced by specialized surface coatings or use of blocking buffers, which would improve efficiency.

**Figure 5.**
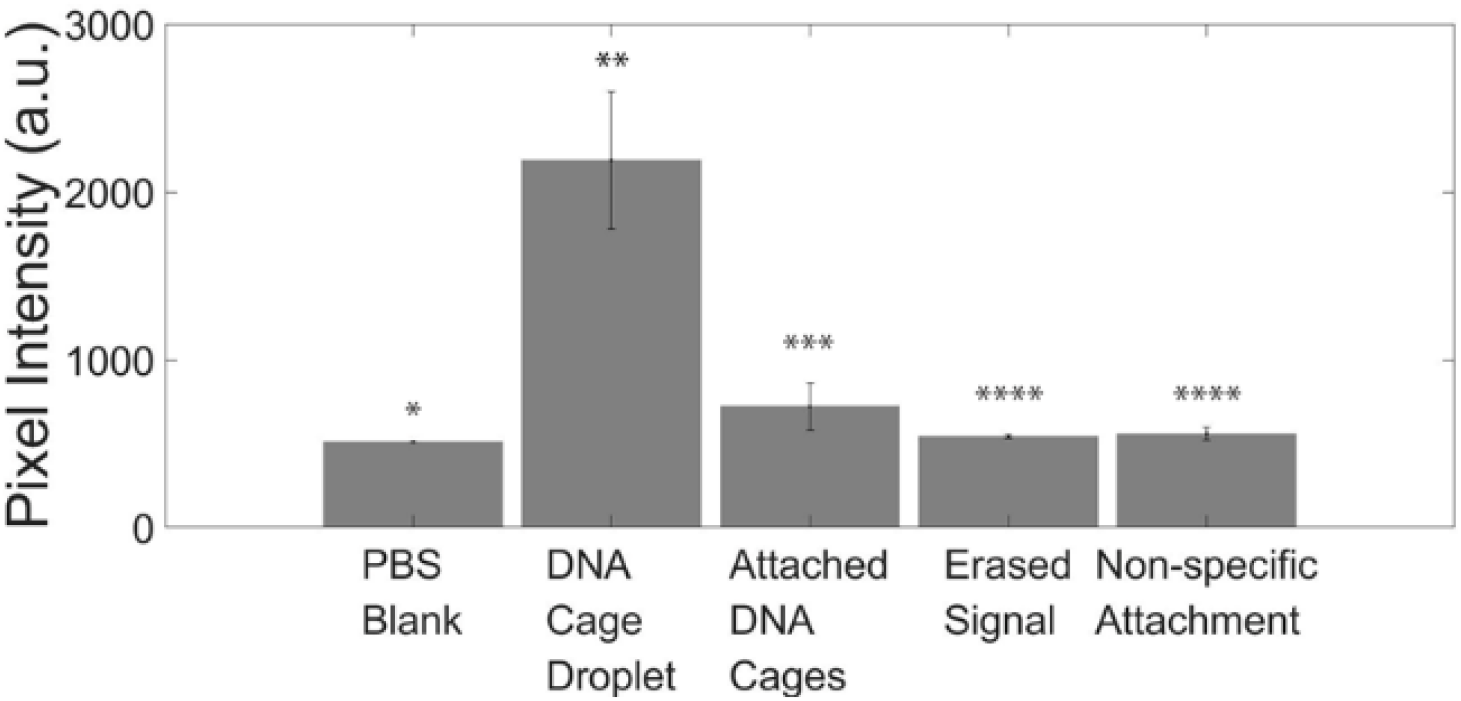
Reversible binding of DNA-caged nanoparticles to ssDNA-coated surfaces. Fluorescent intensity was determined by measuring average pixel intensity from microscope images shown in Supplementary Figure 7. Differing numbers of asterisks indicate statistical differences in pixel intensity between samples.

### Reversible DNA-caged nanoparticle binding in vitro for erasable immunocytochemistry

DNA-modified nanoparticles are often employed in complex intracellular or tissue environments, for example when used as gene therapy or immunotherapy vehicles[1]. To investigate functionality of DNA-caged nanoparticles in cells, an erasable immunocytochemistry model[21] was employed in which coumarin-6 DNA caged-micelles were used to label intracellular actin repeatedly over two cycles (Figure 1B). Primary antibodies were conjugated with ssDNA displaying partial complementarity to DNA cage handles; DNA cages served as the secondary reagent, whose fluorescence was used to identify antigens. DNA-caged nanoparticles were removed from primary ssDNA-antibodies by adding erase DNA with greater complementarity to the handle. Thus, primary ssDNA-antibodies remained in place after erase, and were available for DNA-caged nanoparticle labelling in a second cycle. This particular configuration could find application in Stochastic Optical Reconstruction Microscopy (STORM)[34], which uses repeated imaging of the same antigen to construct super-resolution images. Primary antibodies and ssDNA-primary antibodies with fluorescent secondary antibodies were employed as a positive control; negative controls included coumarin-6 DNA caged nanoparticles without primary antibodies.

DNA-caged nanoparticles displayed similar fluorescence signal (not statistically different) and intracellular distribution to positive controls with unmodified and ssDNA-modified primary antibodies and Alexa Fluor 568 secondary antibodies (Figure 6, Supplementary Figure 8). Negative controls without primary antibodies displayed a signal increase of ∼60% from the blank, which was statistically lower than experimental intensity and representing only 3% of the positive signal. DNA-caged nanoparticles were detached from primary antibodies by addition of erase DNA with erase depths of up to 70% observed. Several factors influenced erase depth. First, if erase was conducted under flow (i.e., no stop-flow incubation steps), only ∼ 20% of signal was erased over the same 15 minute observation period. This is likely a result of poor diffusion combined with convective flow fields. Permitting 15 minutes of stop-flow incubation and adding a PBS washing step increased erase depth to ∼40-50%. Addition of a second identical erase step increased erase depth to ∼ 60-70%. This suggests room for optimization of the erase process, such as altering DNA sequences, concentrations, and reaction times. Also, unlike in solution-based (Figure 4) or surface (Fig 5ure) studies, an increase in residual signal was observed between cycle 1 and 2 that could not be attributed to non-specific binding. This suggests greater difficulty performing strand displacement reactions inside cells, likely reflecting diffusional limitations. Efficiency was increased by washing and additional erase steps, indicating that viscous flow may be necessary to remove erased ssDNA from targets.

**Figure 6.**
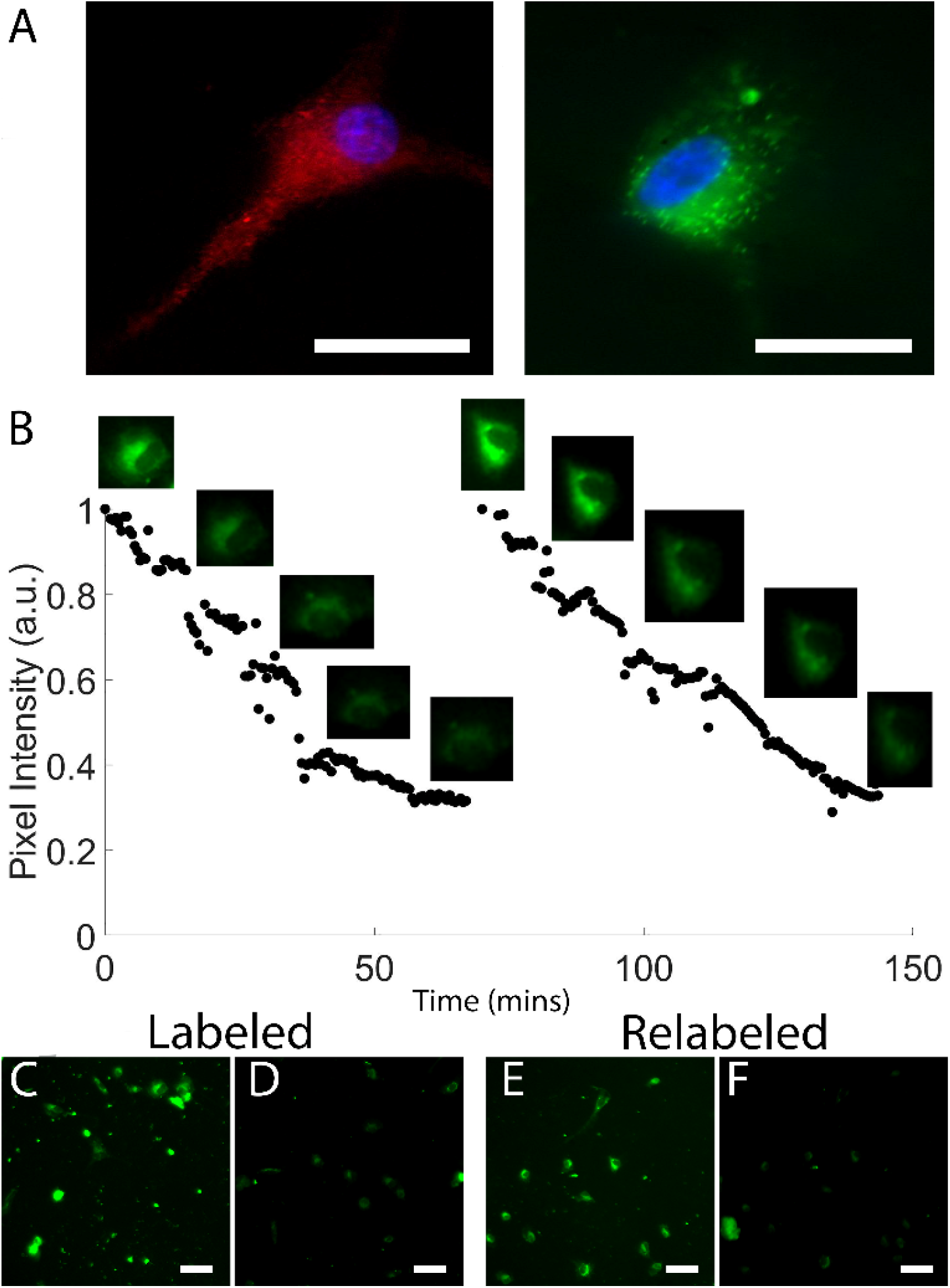
Fluorescent labelling and erasing of β-actin in fixed U87 cells. (A) High resolution images of U87 cells labeled with ssDNA-primary antibodies and (left) control Alexa Fluor 568 secondary antibodies or (right) experimental DNA-caged nanoparticles. Scale Bar = 100 nm. (B) Normalized (to initial) fluorescent intensity of cells labeled with DNA-caged nanoparticles during erase and washing steps imaged under time-lapse. Insets show example images of a single cell at the selected time points. (C-F) Widefield fluorescent images of DNA-caged nanoparticle labeled U87 cells (C) at the start of cycle 1 erase, (D) at the end of cycle 1 final wash, (E) at the start of cycle 2 erase, and (F) at the end of cycle 2 final wash. Scale Bar = 100 nm.

## Discussion

Electrostatic DNA caging technologies offer many potential advantages over traditional bioconjugation schemes[9] that can potentially diminish nanoparticle properties[12, 35] and can have limited yields[11]. DNA cages offer the potential to precisely control ssDNA handle density and position on the nanoparticle surface. Although the approach demonstrated here relied on DNA tiles with incomplete interlocking with stochastic handle presentation, schemes based on interlocking icosahedra[19] or other programmable structures[17, 36] can easily be envisioned. However, unlike other caging strategies, the electrostatic DNA caging approach does not require *a priori* ssDNA modification[37] or use thiolated strands only compatible with a narrow range of nanoparticle types (e.g., gold nanoparticles)[19, 38]. The electrostatic DNA caging approach has broad applicability, requiring only positively charged nanoparticle surfaces. Modifications to cage designs would be required to accommodate nanoparticles of different sizes, and given that the DNA duplex is ∼ 2 nm in diameter there is likely a fundamental size limit. Thus, the technology may not be appropriate for the smallest nanoparticles, such as carbon dots[39].

These data also establish the functionality of ssDNA handles extending from DNA cages in a variety of contexts. Solution-based testing suggests potential compatibility with DNA capture probes, aggregation-based biosensors, and flow cytometry applications. Solid-liquid interface testing establishes relevance of this approach for DNA microarrays. Testing in an intracellular erasable labelling scheme indicates that DNA-caged nanoparticles can function in complex biological environments. Although these experiments repeatedly labelled the same antigen, a configuration amenable to STORM imaging[34], use of different antibodies could be employed for multiplexed imaging. Erasable fluorescence labelling has been reported in several contexts[21, 40-42], including development into commercial systems (e.g., GE MxIF, Nanostring GeoMx®, Akoya Opal Polaris). Erasable labelling enhances multiplexing, enabling use of spectrally overlapping fluorophores. However, most existing approaches rely on UV photobleaching[40] or harsh chemicals[41] to degrade fluorophore signals. In contrast, this approach uses gentle DNA strand invasion to erase label signals. Whereas DNA-based erasable labelling has been previously reported[21], those methods use individual fluorescently labelled ssDNA strands. There is no amplification of fluorescent signal, such as that achieved by micelle encapsulation of multiple dyes, and dye molecules are not protected from the external environment, which can increase photobleaching[43]. Thus, DNA-caged nanoparticles and strand invasion erase may provide an attractive alternative for multiplexed erasable labelling platforms.

## Conclusions

Here, we demonstrate a new method of modifying nanoparticles with DNA based on electrostatic attraction. DNA has a negatively charged backbone that permits electrostatic adsorption of DNA nanostructures on positively charged nanoparticle surfaces. Using Y-shaped DNA tiles stabilized by hybridized interlocks, we created and characterized DNA caged-nanoparticles demonstrating tile adsorption, nanoparticle surface saturation, and partial tile interlocking. We showed that the electrostatic DNA caging approach is versatile, applying it to polymer micelles, polystyrene beads, gold nanoparticles and SPIONs with sizes ranging from 5 to 20 nm. DNA cages formed from tile interlocking were stable in biological buffers and solutions. The DNA cage design provides ssDNA handles that can be used for reversible attachment and binding, which we validated in the solution phase, at solid-liquid interfaces, and in fixed cell cultures. The latter application shows proof-of-concept for DNA-caged nanoparticles in erasable multiplexed immunocytochemistry with the advantages of gentle erase via DNA dehybridization, signal amplification by dye aggregation in polymer micelles, and dye protection from photobleaching by micelle coatings. Thus, the electrostatic DNA caging approach offers a facile method to modify nanoparticles with DNA, providing several advantages to traditional conjugation approaches in that it does not disturb nanoparticle surfaces, is broadly applicable, and permits control of DNA density and pattering. Given these advantages, electrostatic DNA caging could enable more rapid integration of nanoparticles in medicine and broaden access to DNA-nanoparticle composite materials.

## Author Contributions

Author contributions are provided using CRediT descriptions for each author indicated by their initials. EJ: Data Curation, Formal Analysis, Investigation, Validation, Visualization, Writing- Original Draft, Writing-Review and Editing; SF: Data Curation, Formal Analysis, Investigation, Validation, Visualization, Writing-Original Draft, Writing- Review and Editing; YC: Data Curation, Investigation, Visualization, Writing- Original Draft, Writing- Review and Editing; AR: Investigation, Methodology, Resources, Writing- Review and Editing; CEC: Methodology, Writing- Review and Editing; MGP: Methodology, Resources, Supervision, Writing- Review and Editing; MNG: Formal Analysis, Methodology, Resources, Software, Validation, Visualization, Writing- Review and Editing; JJO: Formal Analysis, Methodology, Resources, Software, Supervision, Writing- Review and Editing; JOW: Conceptualization, Funding Acquisition, Methodology, Project Administration, Resources, Supervision, Writing- Original Draft, Writing- Review and Editing.

## Acknowledgements

The authors gratefully acknowledge funding from the Department of Energy Basic Energy Sciences DE-SC0017270 for materials design and realization and the National Science Foundation DBI-1555470 for biological testing and imaging. We acknowledge the Pelotonia Cancer Research Foundation and the Center for Cancer Engineering–Curing Cancer Through Research in Engineering and Sciences (CCE-CURES) for support of Silvio de Araujo Fernandes-Junior. The authors also acknowledge a generous donation from the Paul Bigley family. This work is dedicated to his memory.

