## Supplementary Material for "DNA-caged Nanoparticles via Electrostatic Self-Assembly"

**Supplementary Methods**

Characterization of micelle nanocomposite size, encapsulation efficiency, and polydispersity

To determine the morphology and size of the polymer nanoparticles, we used transmission electron microscopy (TEM). TEM grids were treated using a PELCO easiGlow™ Glow Discharge Cleaning System prior to nanoparticle deposition. Then, a 12.5 μL sample droplet was pipetted onto a silicone pad over which the TEM grid was inverted for 5 minutes. Excess sample was then slowly wicked away using filter paper. Negative staining was performed using 1% uranyl acetate dissolved in distilled water. TEM images were collected using an FEI Tecnai G2 Bio Twin TEM at 80kV.

Encapsulation efficiency was measured using ultraviolet-visible (UV-Vis, GENESYS 6, Thermo-Fisher) measured against a standard curve. Coumarin-6 micelles were isolated using centrifugal filtration (regenerated cellulose 30 kDa NMWL Amicon Ultra-15, Millipore Sigma cat. no. UFC903024) at 4000 rpm for 15 minutes with 3 washes using DI water. Then, collected samples were imaged in UV-Vis and compared to the signal for the coumarin-6 containing initial solution. The ratio of these two signals provides the encapsulation efficiency.

**Supplementary Table 1. DNA tile sequences used**

| G1-sticky-handle | AAAAATTTCGACGTTACATGCACCTCGCTCGAGCCAGTGAGGACGGAAGTTTGTCGTAGCATCGCACC |
| --- | --- |
| G2-sticky-handle | AAAAATTTCGACGTTACATGCACCTCGCTCGAGC CAACCACGCCTGTCCA TT ACTTCCGTCCTCACTG |
| G3-sticky-handle | AAAAATTTCGACGTTACATGCACCTCGCTCGAGC GGTGCGATGCTACGAC TT TGGACAGGCGTGGTTG |
| 9 bp compliment | TAAATTGAGGATTATCAAACATGTAACG/3Cy5Sp/ |
| 12 bp compliment | ATTGAGGATTATCAAAGAGGTGCATGTA |
| 15 bp compliment | GATTATCAAAGAGGTGCATGTAACG/3Cy5Sp/ |
| 26 bp (full) compliment | GAGGTGCATGTAACGTCGAAATTTTT |
| Slide | GATTATCAAAGAGGTGCATGTAACGTCG/3ThioMC3-D/ |
| Antibody | /5Cy5/GATTATCAAAGAGGTGCATGTAACGTCG/3AmMO/ |

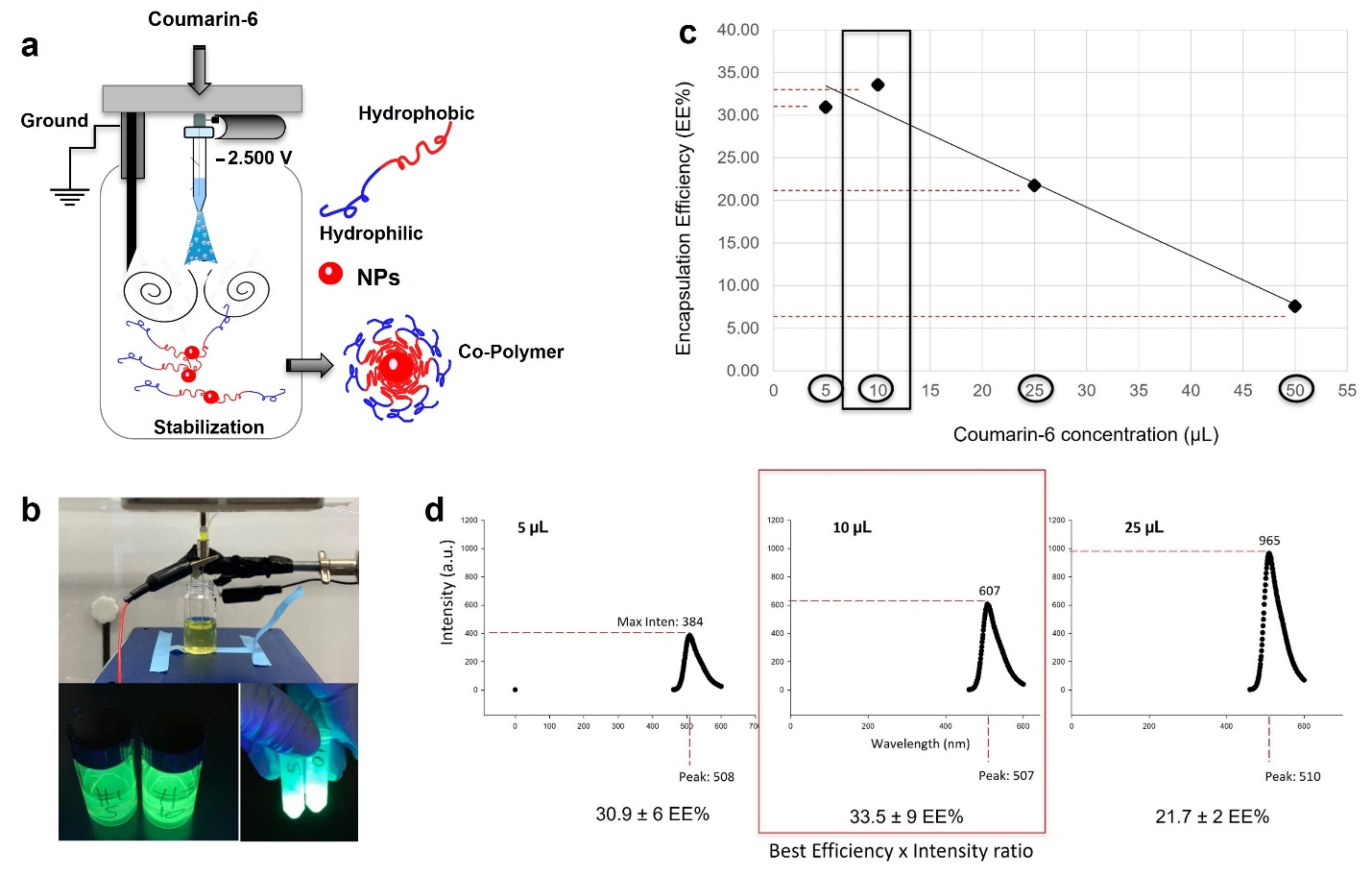

**Supplementary Figure 1:** (a) Schematic of the electrohydrodynamic mixing high voltage nanoprecipitation of amphiphilic DSPE-PEG polymers loaded with fluorescent coumarin-6 dye (C6). (b) (Top) Actual set-up and (bottom) example coumarin-6 micelles (bottom, left) before and (bottom, right) after purification. (c) Coumarin-6 UV-Vis calibration curve showing linear range. (d) Encapsulation efficiencies of coumarin-6 micelles with different amounts of coumarin-6 added. Optimal encapsulatio efficiency occured at 10 μL.

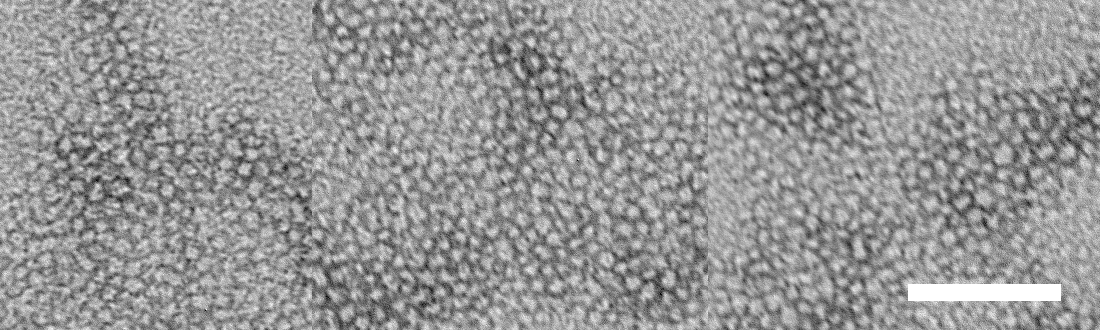

**Supplementary Figure 2.** TEM images of coumarin-6 micelles. Scale bar = 100 nm.

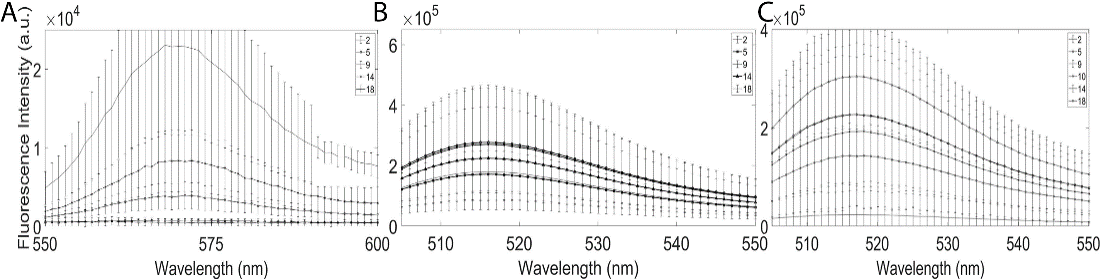

**Supplementary Figure 3.** Surface quenching, surface saturation, and tile interlocking on the surface of DSPE-PEG nanoparticles as a function of DNA: polymer molar ratio (upper right) with error bars. Figures without error bars are shown in Figure 3.

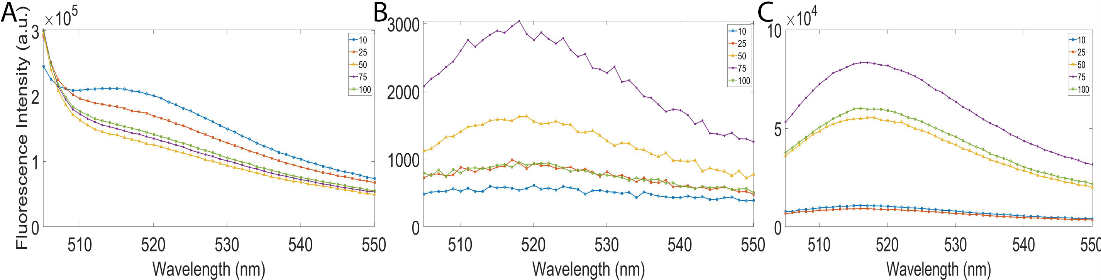

**Supplementary Figure 4.** Electrostatic adsorption of DNA tiles modified with FAM-6 dye to aminated A) 20 nm polystyrene (PS) beads, B) 20 nm gold nanoparticles (AuNPs), and C) 5 nm super paramagnetic iron oxide nanoparticles (SPIONs) as a function of nanoparticle:  DNA volume ratio (upper right).

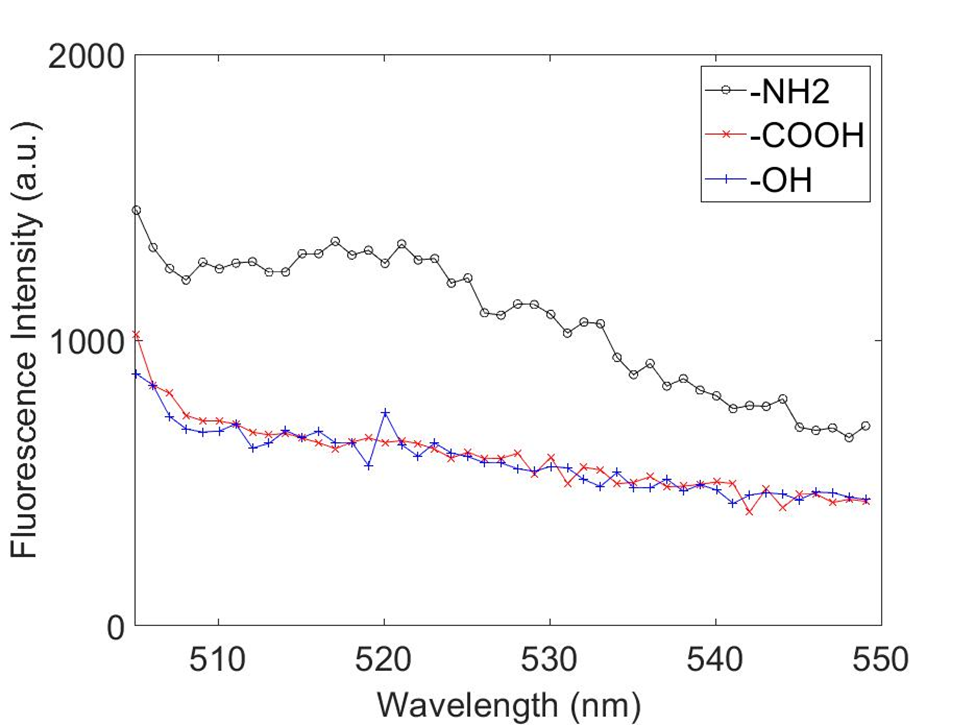

**Supplementary Figure 5.** Electrostatic adsorption of DNA tiles modified with FAM-6 to 20 nm polystyrene (PS) beads with different surface modifications (upper right) at a fixed nanoparticle: DNA volume of ratio of 10:1.

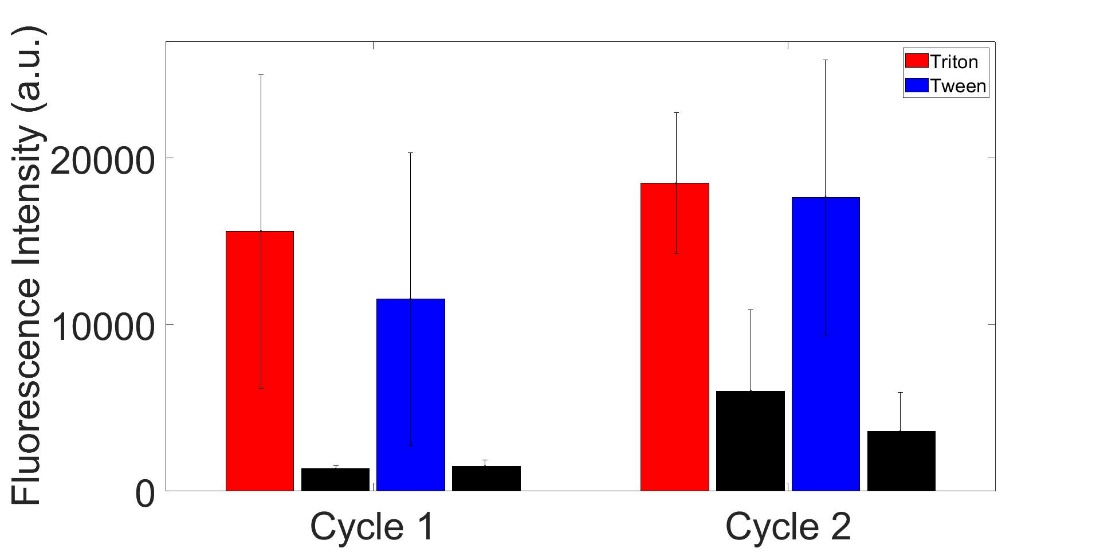

**Supplementary Figure 6.** DNA caged nanoparticle ssDNA handle functionality as determined by strand invasion in the presence of 0.25% Triton X100 or 0.25% Tween-20 solutions. Fluorescent, Cy5-ssDNA sequences with partial complementarity to handles were added and subsequently detached via strand invasion using non-fluorescent ssDNA sequences with greater handle complementarity. Fluorescent intensity of DNA-caged nanoparticles after Cy5-ssDNA binding and erase was measured over two cycles.

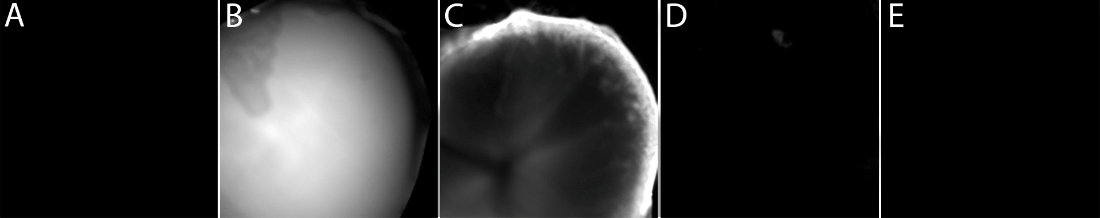

**Supplementary Figure 7.** Fluorescent microscope images of DNA caged nanoparticles on slides: A) PBS blank before DNA-caged nanoparticle addition, B) after addition of a droplet of DNA-caged nanoparticles, C) DNA-caged nanoparticle signal following droplet removal and washing with PBS three times (attached), D) residual DNA-caged nanoparticle signal after strand displacement and washing with PBS three times (erased), and E) non-specific attachment of DNA cages to slides without ssDNA binding targets.

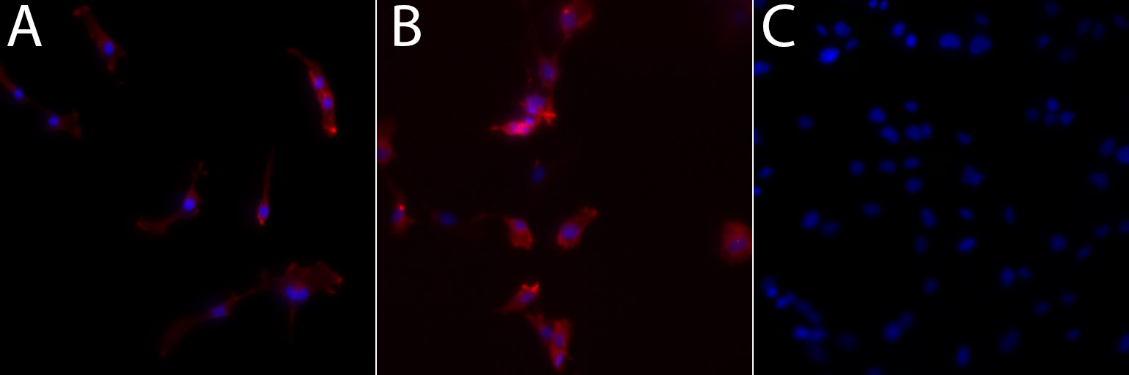

**Supplementary Figure 8.** Fluorescent microscope images of cell labeling controls: A) Cells labeled with primary antibodies and Alexa Fluor 568 secondary antibodies, B) Cells labeled with ssDNA primary antibodies and Alexa Fluor 568 secondary antibodies, and C) Negative control with no primary antibodies and DNA caged nanoparticles added.
